## Supplementary manuscript for "Inferring residue level hydrogen deuterium exchange with ReX"

### 1 Additional Benchmarking Results

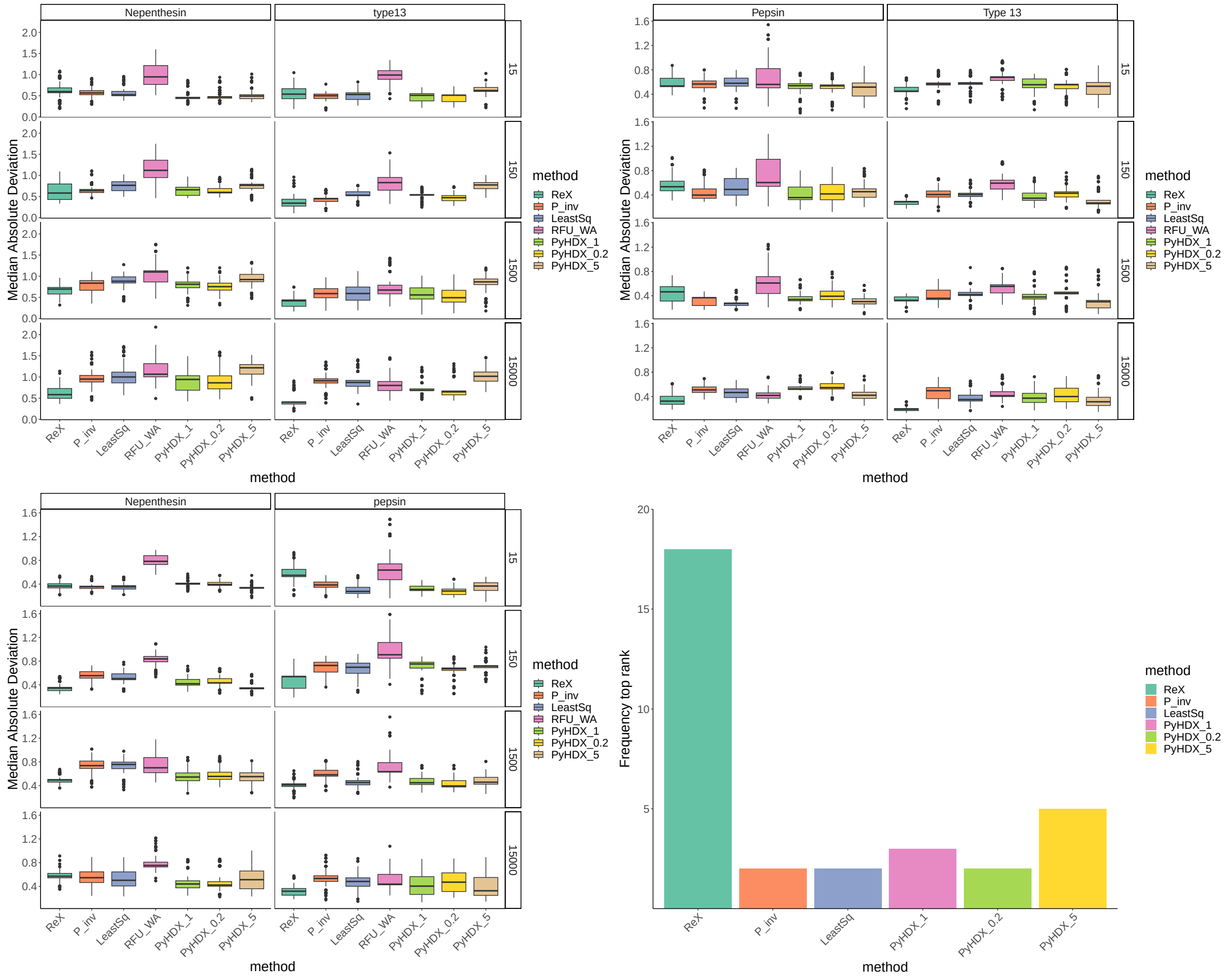


Figure S1: **Additional Benchmarking results.** Benchmarking results comparing different residue resolved approaches. Boxpots are in the style of Tukey presenting approximate 95% confidence intervals. The boxplots represent bootstrap distributions of the Median Absolute Deviation (MAD) of the predicted and observed values. (Upper Left) trained on Pepsin (IUpper Right) Trained on Nepenthesin (downsampled to 40 peptides) (Lower Left) trained on aspergillus pepsin type XIII. (Bottom Right) Summary of results.

#### 2 Error Distributions

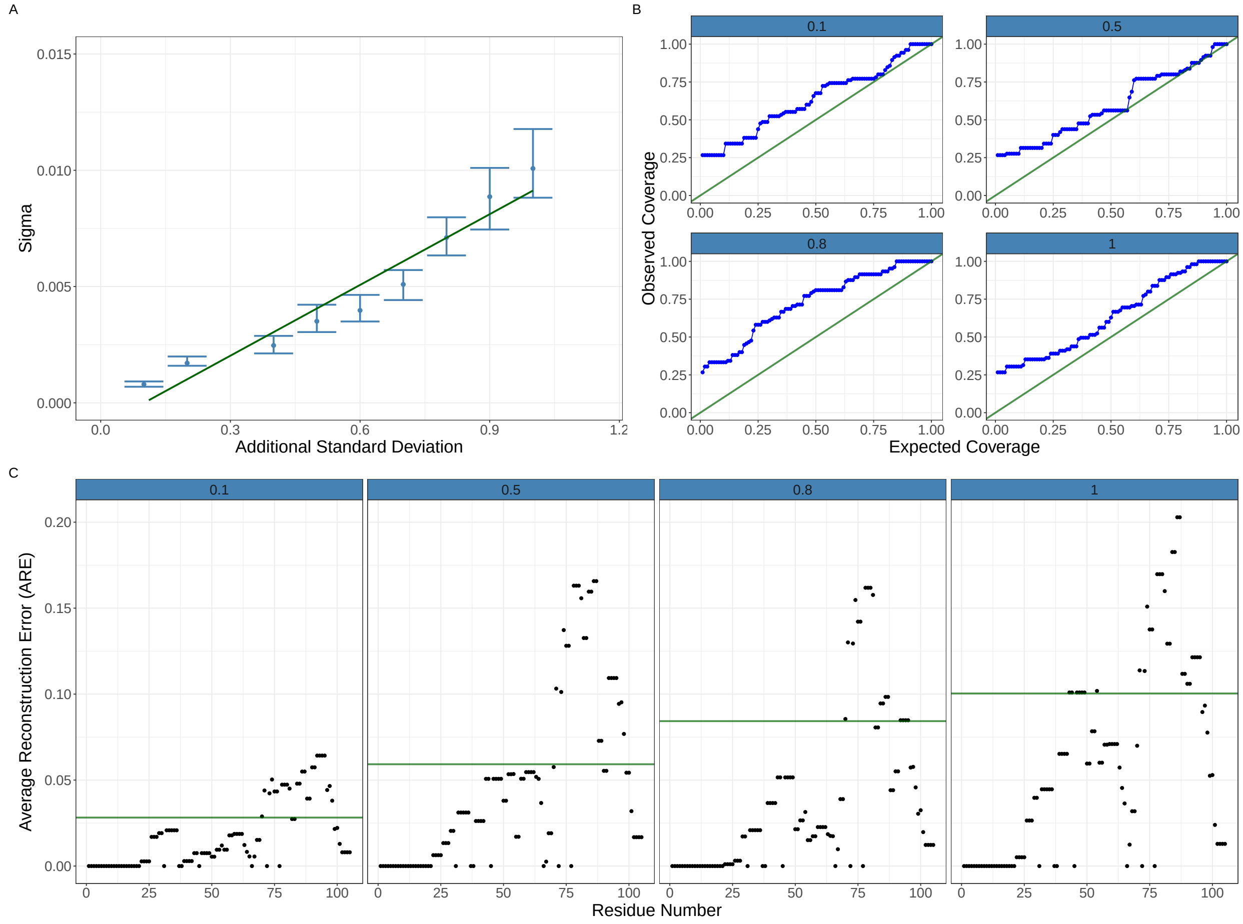


Figure S2: **Error distribution Figure.** Plots representing how adding additional standard deviations effect the quality measures. (A) The added additional standard deviation and the inferred Sigma value. Equitailed 95% credible interval reported. Straight line fitted to data demonstrated close to linear increasing relationship between the additional standard deviation and the inferred Sigma. (B) Coverage plots showing that credible interval constructed using Sigma overestimate the variability in the data and are hence conservative. Plots are faceted by additional standard deviation. (C) Average Reconstruction Error (ARE) plotted against Residue Number. The dark green line is the inferred median value of the square root of Sigma and the plots are faceted by additional standard deviation. ARE and Sigma are faithful to each other and the additional standard deviation.

### 3 Structural annotations distributions

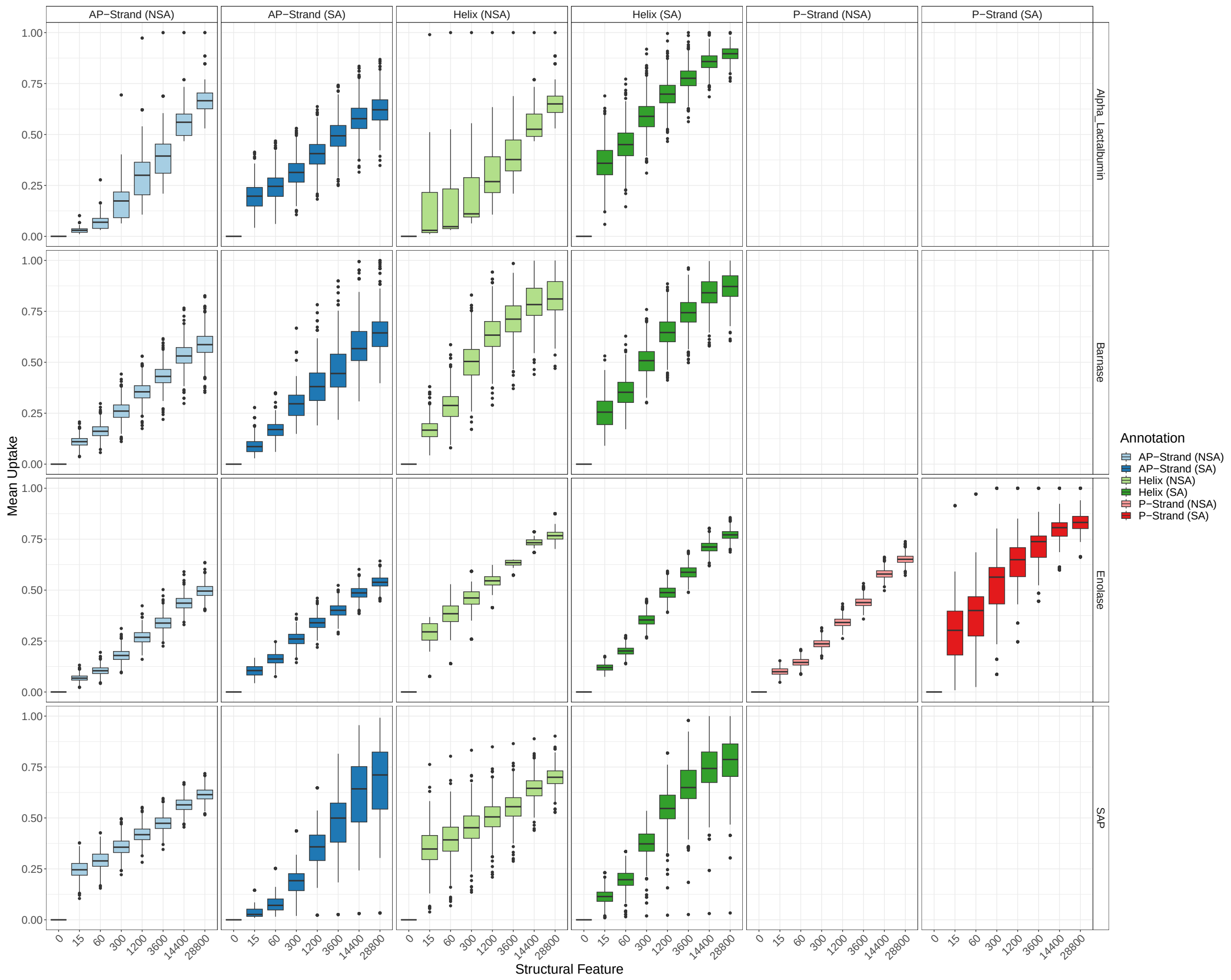


Figure S3: **Structural Features Boxplots.** Box plots corresponding to main Fig. 4B. The distribution are over random subsamples, are x-faceted by structural feature and y-faceted by protein.

### 4 Differential HDX-MS simulations Meta-Analysis

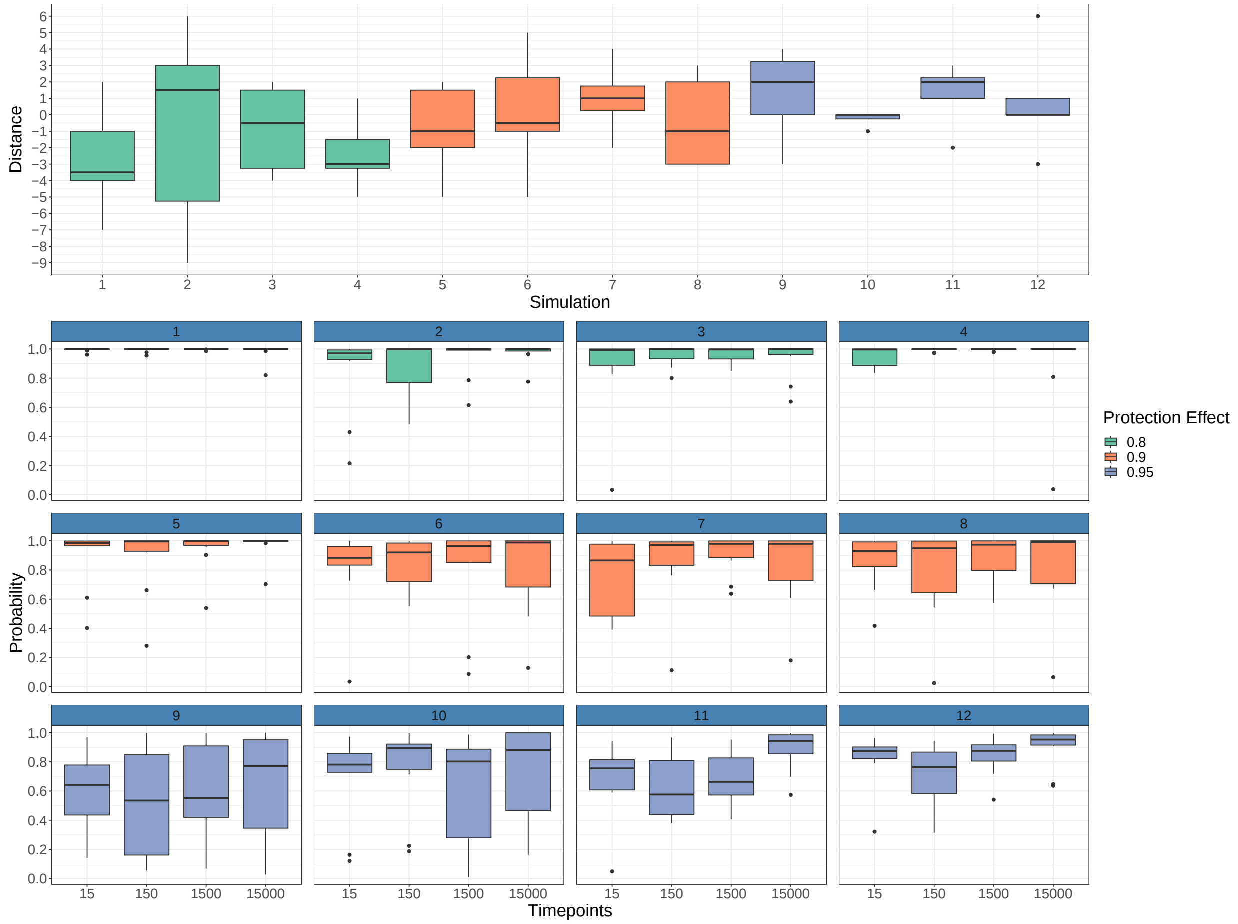


Figure S4: **Meta-analysis of simulations.** Complete results of simulation study. Simulations are index one through twelve with each simulation repeated over ten random seeds. Upper plot show distance from residue of maximal probability to residue that was selected for perturbation. Lower plot is faceted by simulation and indexed by time points. The boxplots show the probability given to residue selected for the simulation. Note that an 80% protection effect is a 20% reduction in deuterium uptake.

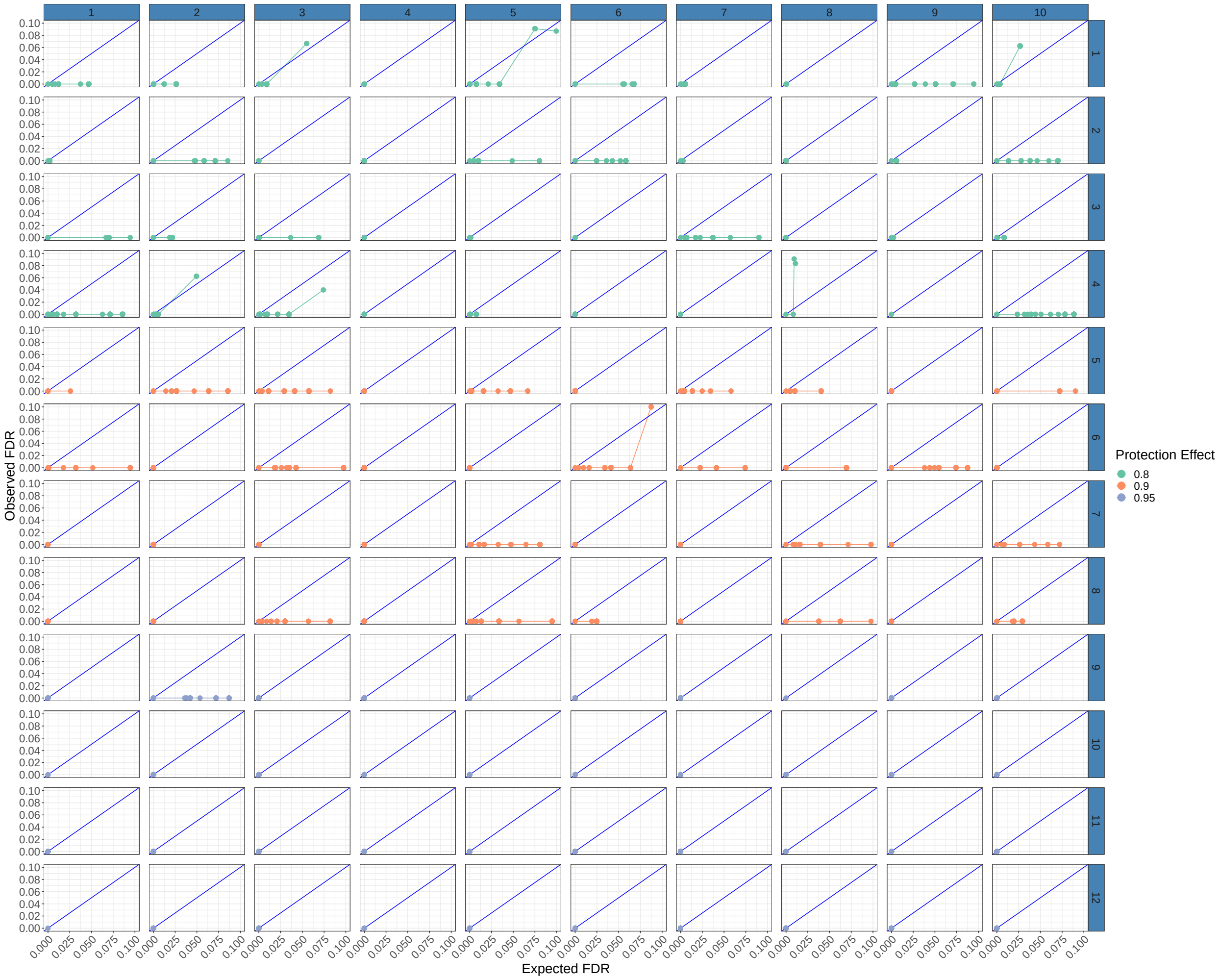


Figure S5: **Calibration plots.** Calibration plots showing expected FDR against observed FDR. Note that the lines are mostly below the y-x axis. The data is x-faceted by random seed and y-faceted by simulation. Note that not all EFDR levels are computable from the simulations.

### 5 Additional LXRα analysis

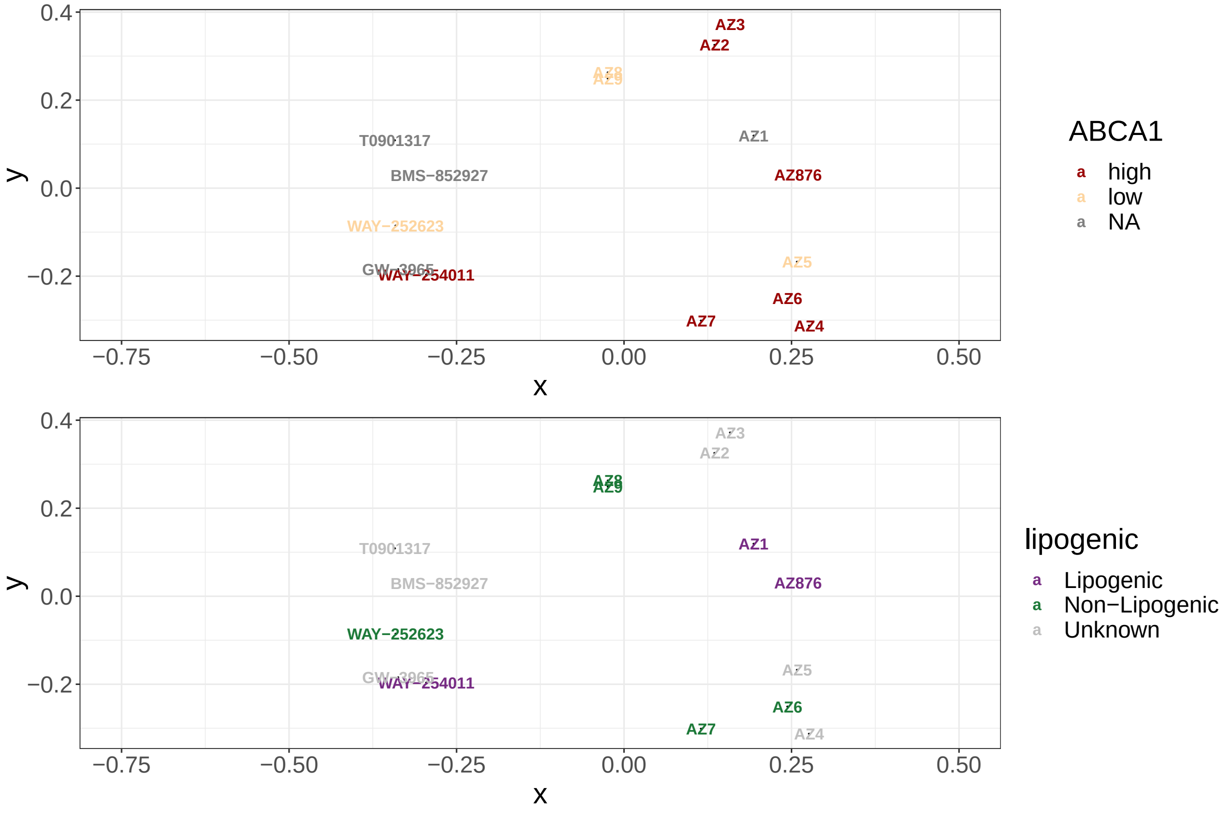


Figure S6: **Multidimensional scaling of chemical similarity** Each figures show a Multidimensional scaling (MDS) scaling plot the separation of the ligands in chemical (Tanimoto) similarity. Neither $x$ nor $y$ components separate in vivo determined functions. Compounds that induce conformationally distinct profiles cluster with similar compounds. Indicating conformation is more closely linked to function that chemical similarity.

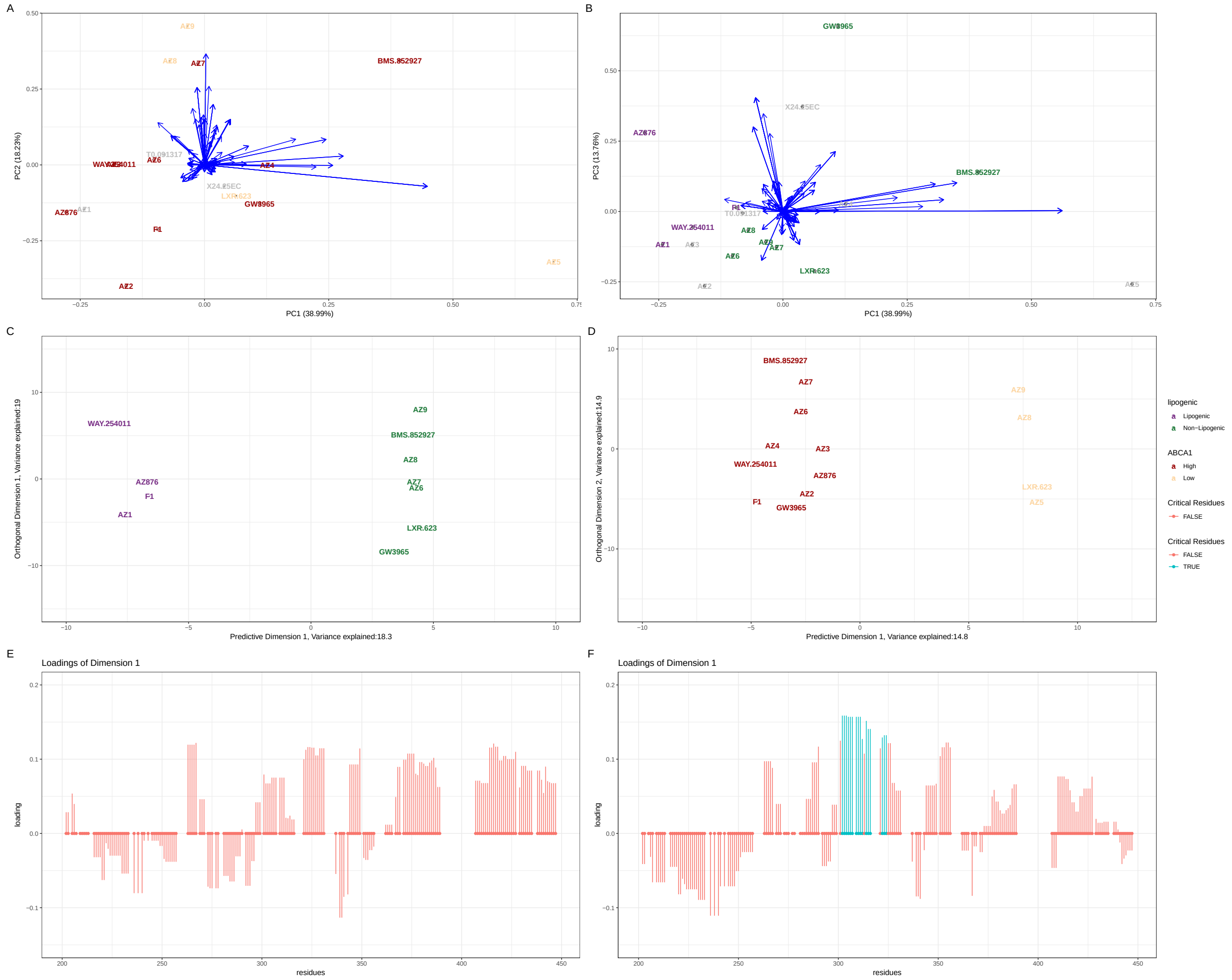

Figure S7: **LXRα in complex with SRC1 CSA.** (A) PCA plots of HDX-MS signatures per ligand. Ligands are coloured by ABCA1 induction and blue arrow represent residue contributions. (B) As with (A) but for PC1 versus PC3. The ligands are coloured by lipogenic annotation. (C) An OPLS-DA plot with lipogenic as the outcome variable (D) An OPLS-DA plot with ABCA1 induction as a outcome variable (E) The loading plot corresponding to the predictive dimension of (C) larger loadings indicate greater contribution. (F) A loading plot corresponding to the predictive dimension of (D).

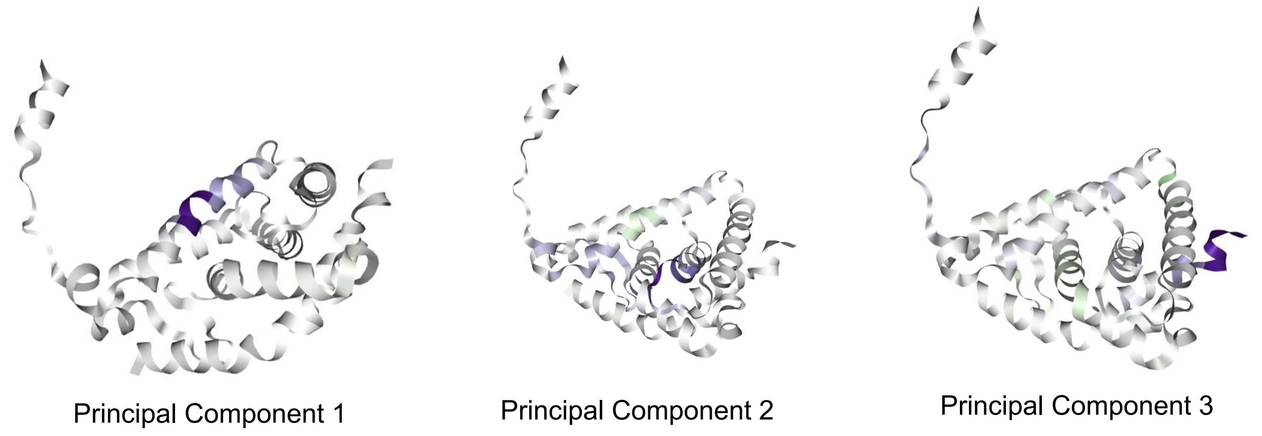


Figure S8: **LXR$\alpha$ in complex with SRC1 CSA PCA loadings.** Loadings corresponding to the annotated principle components from Fig. S7 A and B. Plotted directly on the structure of LXR$\alpha$

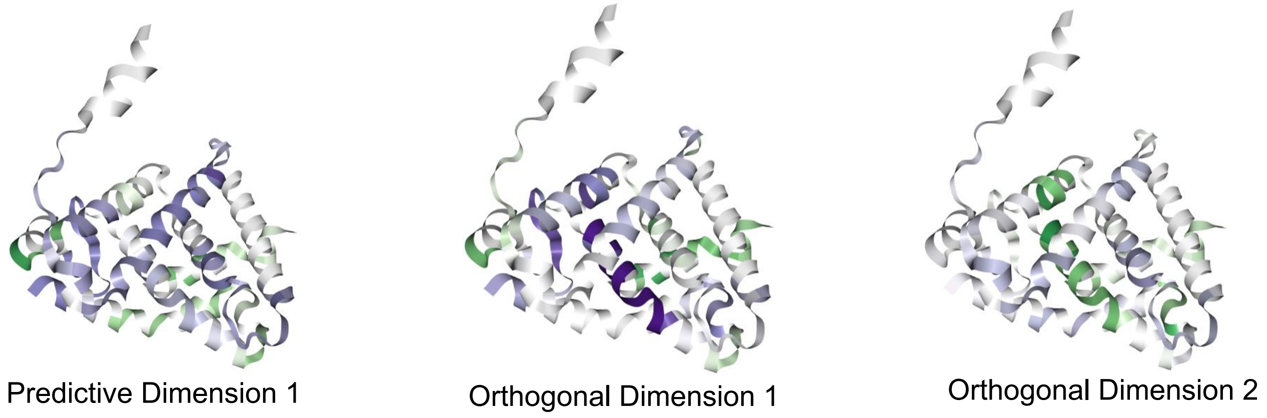


Figure S9: **LXR$\alpha$ in complex with SRC1 CSA OPLS-DA loadings.** Loadings corresponding to the annotated OPLS-DA from Fig. S7 A and B. Plotted directly on the structure of LXR$\alpha$

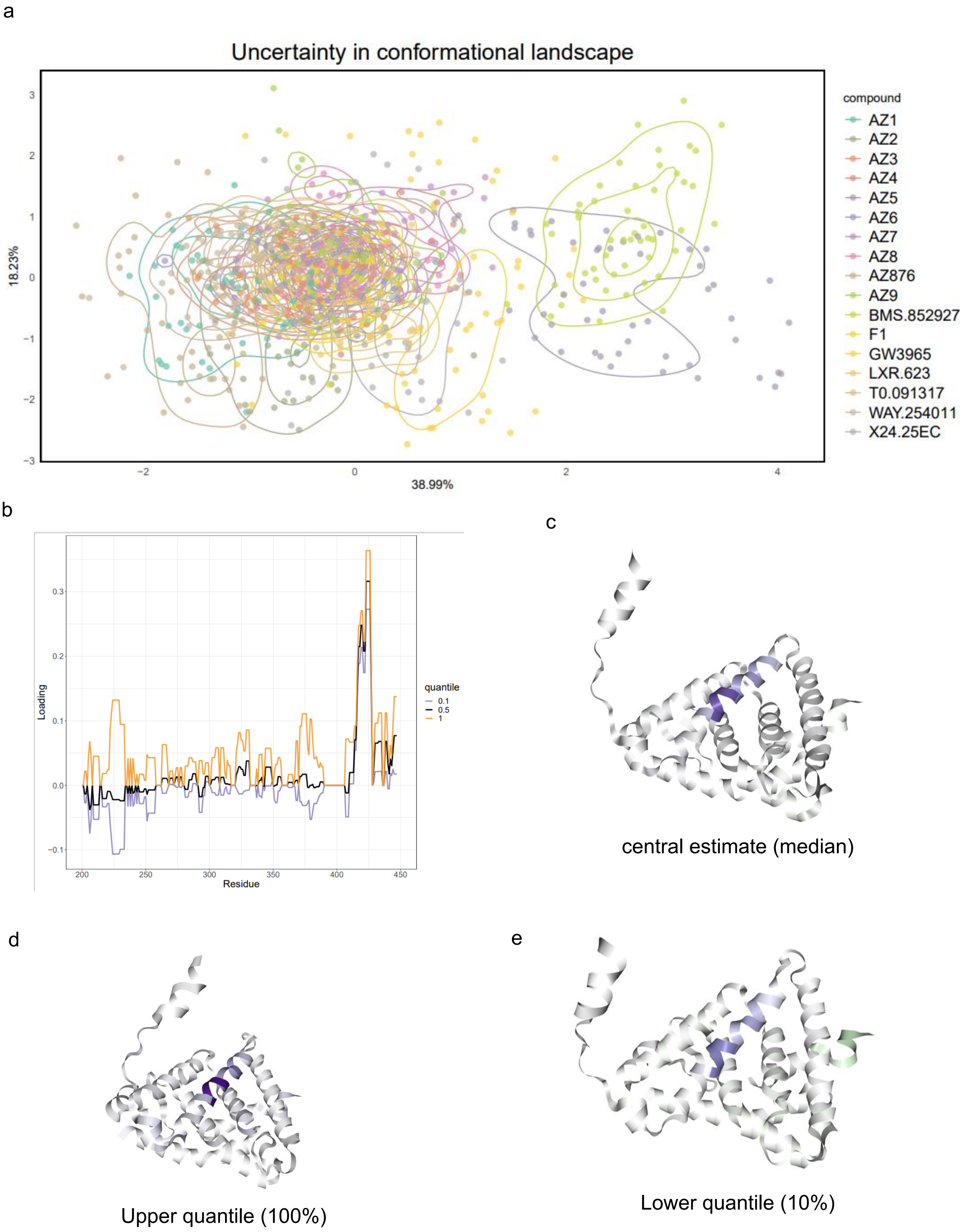


Figure S10: **LXR$\alpha$ in complex with SRC1 CSA Uncertainty Quantification.** (A) A PCA (PC1 vs PC2) plot with uncertainty contours representing the posterior distribution of the locations of molecules in PCA coordinates. (B) The corresponding quantiles in the loadings of the principal components (C) The loading plotted directly on the protein structure corresponding to the mean estimate of the loadings. (D) As with (C) but the upper quantile of the loading (E) as with (D) but with the lower quantile of the loadings.

### 6 BRD protein alignments

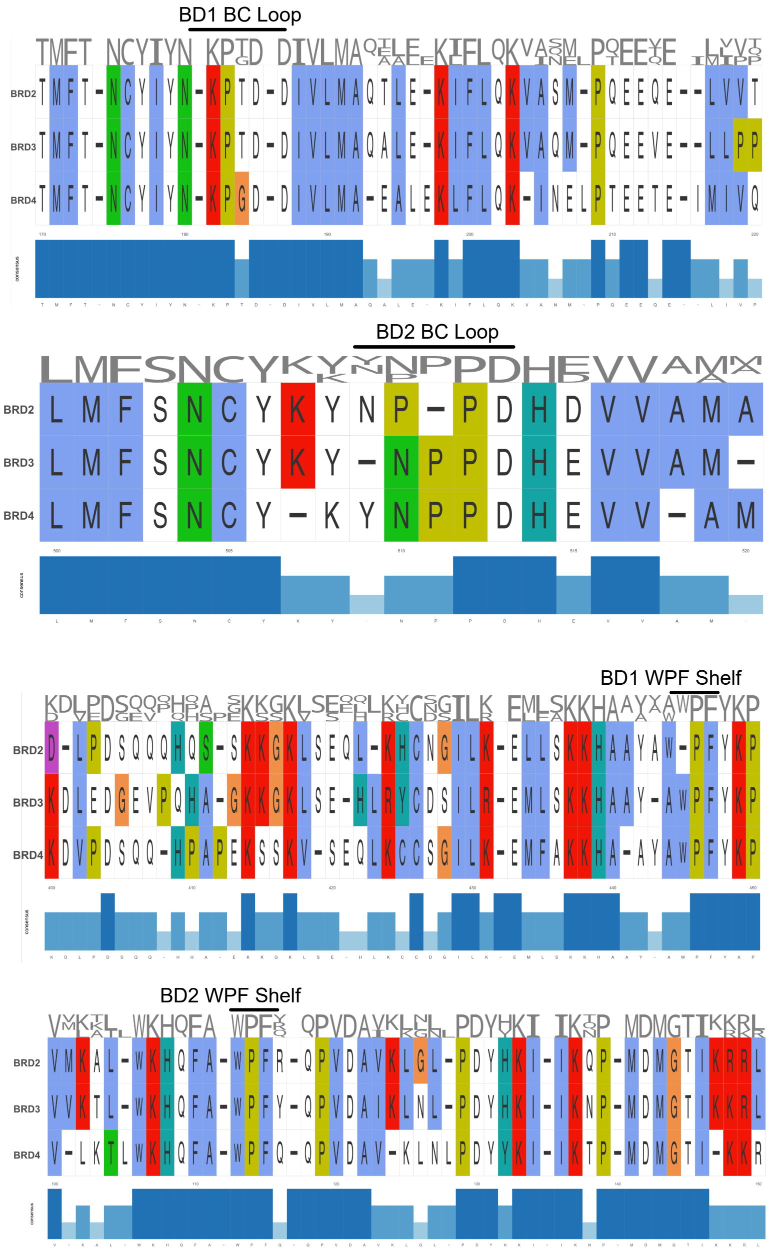


Figure S11: **BRD 2,3 and 4 sequence alignment** Sequence alignments (Clustal) for BRD2,3 and 4 focusing on the BC loop and WPF shelf.

### 7 Structure Evaluation from HDX

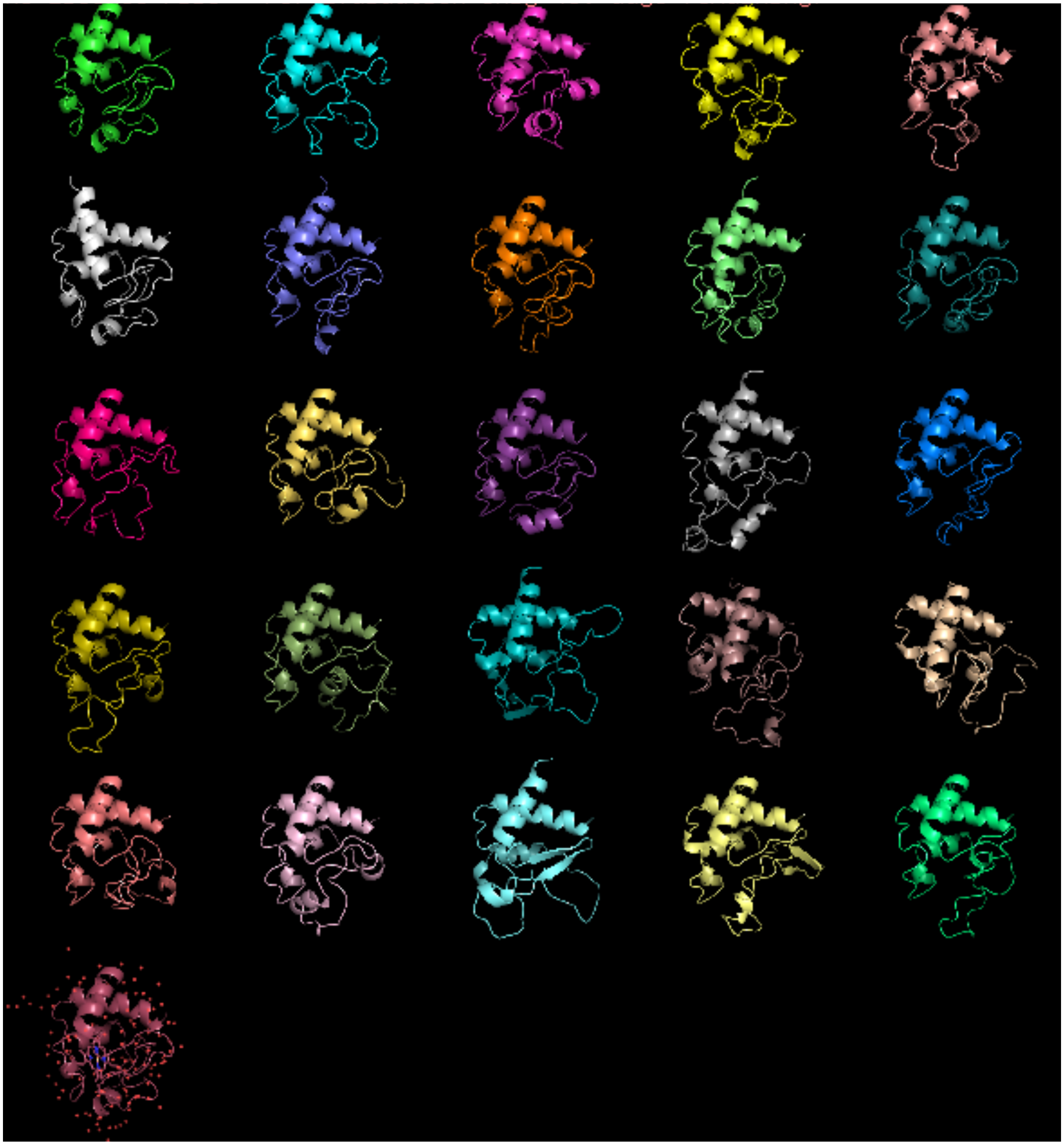

Figure S12: **AlphaFold models of Cytochrome C.** AlphaFold (AF) model of Cytochrome C generated using multiple sequence alignment (MSA) subsampling[1]. The models are wrapped row-wise in order and the crystal structure is given for reference. Note that heme is in the crystal structure (PDB: 1HRC) and this co-factor is not used to generate the AlphaFold structures. Cytochrome C is also in AF trainings set.

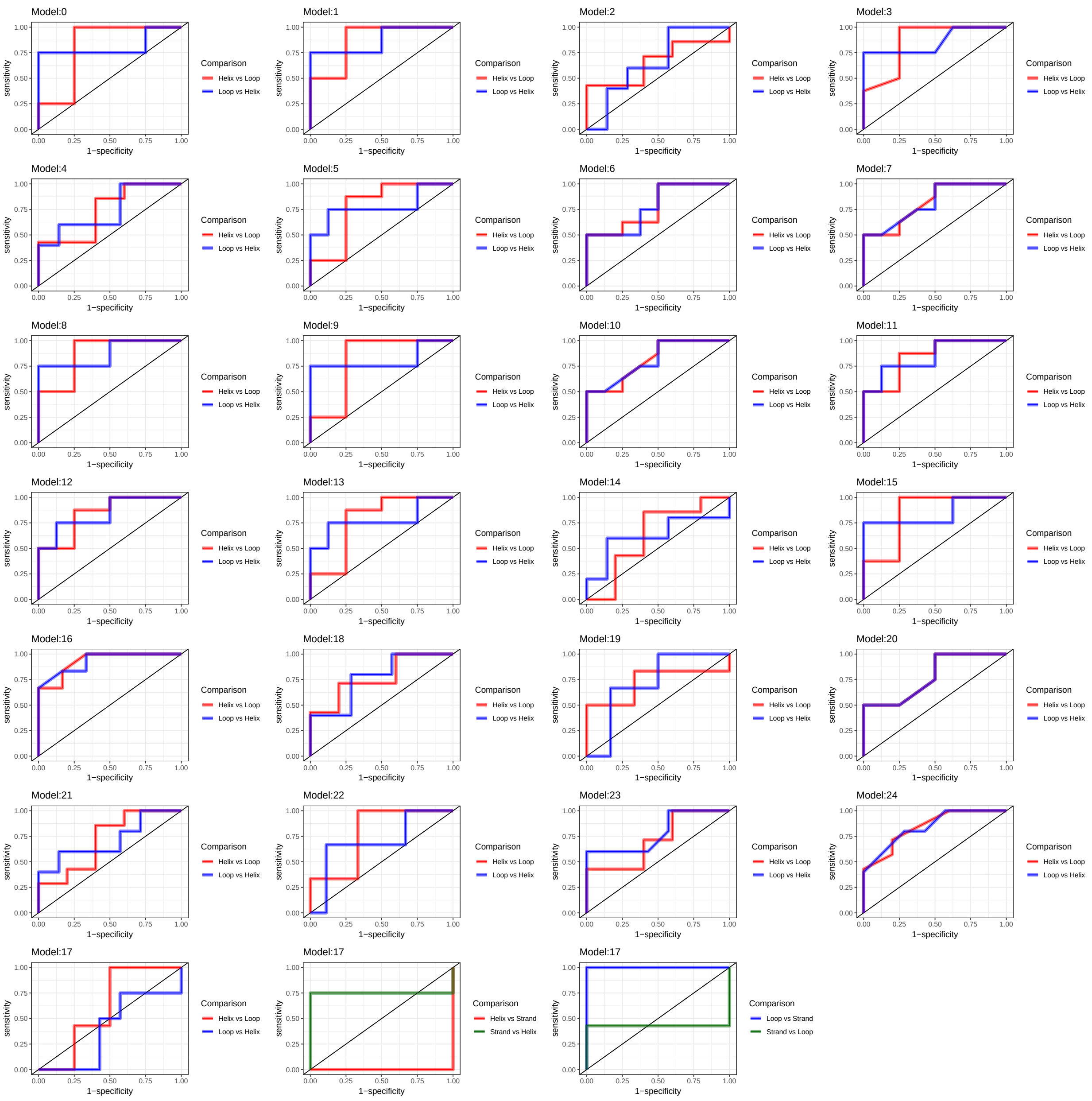


Figure S13: **ROC curves for Secondary Structure prediction.** ROC curves showing predictive performance of HDX feature in predicted secondary structure elements. Curves are shown for both prediction directions. A holdout dataset of 20% of residues is used for each predicted model.

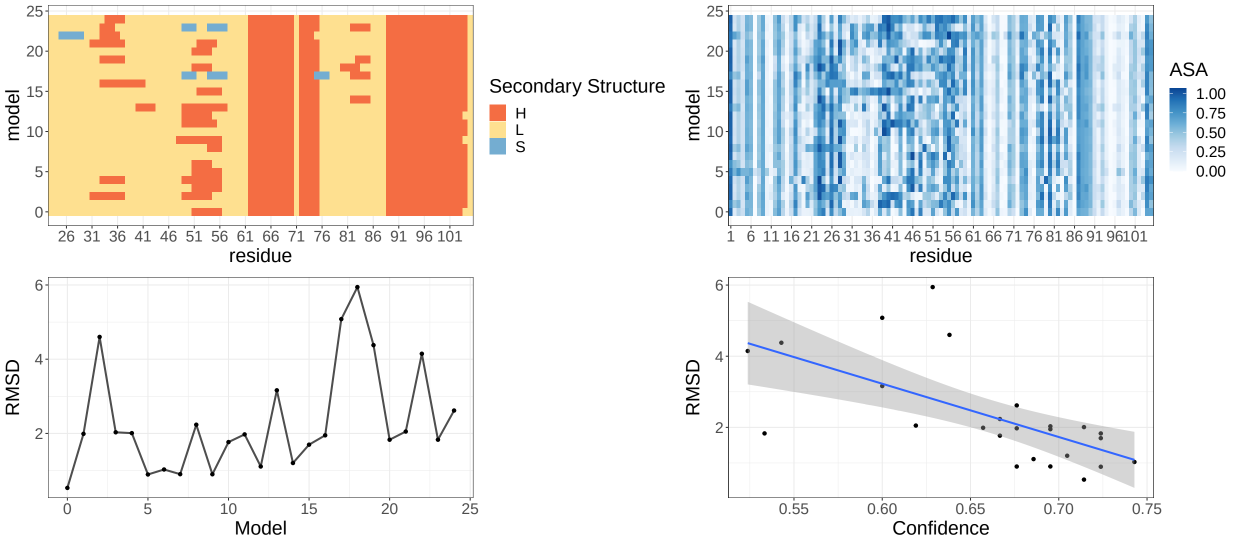


Figure S14: **Evaluating confidence in AlphaFold models.**(Top left) Secondary Structure annotations for each model. (Top right) Accessible solvent area for each model. (Bottom left) RMSD to crystal structure of each model. (Bottom right) RMSD as a function of random forest derived confidence. Higher confidence leads to lower RMSD. Linear model shown with standard error confidence band.

### 8 Supplementary methods

#### 8.1 AF-Cluster

AlphaFold conformation were generated as described in Wayment-Steele et al. [[1]] with version dated Jan 2023 from the Google collab notebook. In particular a multiple sequence alignment (MSA) of Cytochrome C was generated and clustered. The method was run with default settings with three recycles and model number set to three. This generated a total of 25 candidate models.

#### 8.2 Secondary structure assignment

The Secondary structure of Cytochrome-C was re-annotated using STRIDE[2].

#### 8.3 Prediction of secondary structure

The Secondary structure was predicted per model using a random forest model with an 80/20 split between training and testing.

#### 8.4 Accessible surface area

Relative ASA was calculated using FreeSASA[3] using the standard spherical probe model.

#### 8.5 Confidence scores

Root mean squared deviation (RMSD) calculations were made between each model to the crystal structure (PDB: 1HRC). Using a random forest model we predicted both secondary structure and accessibility (ASA $> 0.25$) with a 80/20 residue train/test split. The prediction probabilities for secondary structure and ASA were obtained from the model and a per residue confidence score was calculated by multiplying these probabilities together. A per model confidence score was obtained by computing the proportion of residue confidence scores above 0.5.
